# Sex differences in donor T cell targeting of host splenocyte subpopulations in acute and chronic murine graft-vs.-host disease: implications for lupus-like autoimmunity

**DOI:** 10.1101/2024.06.07.595177

**Authors:** Kateryna Soloviova, Charles S. Via

**Author notes:** Address correspondence to: Charles S. Via M.D., Department of Pathology, Room 3B100, 4301 Jones Bridge Road, Uniformed Services University of Health Sciences, Bethesda, MD 20814. The opinions expressed herein are those of the authors, and are not necessarily representative of those of the Uniformed Services University of the Health Sciences (USUHS), the Department of Defense (DOD); or, the United States Army, Navy, or Air Force.

## Abstract

This study sought to compare in vivo sex differences in either a Th1-dominant CTL response or a Tfh-mediated lupus-like antibody response using the parent-into F1 murine model of acute or chronic GVHD respectively. In acute GVHD we observed no significant sex differences in the hierarchy of donor CD8 CTL elimination of splenocyte subsets. B cells were the most sensitive to elimination in both sexes; however, the male response was significantly stronger. Sex differences in chronic GVHD were more widespread; females exhibited significantly greater numbers of total splenocytes and host CD4 Tfh cells, B cells and CD8 T cells consistent with reports of greater female autoantibody production in this model. The more potent male CTL response in acute GVHD conflicts with reports of greater female CTL responses following infections or vaccines and may reflect the absence of exogenous innate immune stimuli in this model.

## 1. Introduction

Systemic lupus erythematosus is multi-system autoimmune disease characterized by a waxing and waning course and significant clinical heterogeneity that together make human mechanistic studies difficult [1]. As a result, animal models have been of enormous value in elucidating important pathogenic mechanisms involved in the disordered immune function characteristic of human lupus. A useful mechanistic model for the study of lupus is the parent-into-F1 (P→F1) model in which a lupus like disease can be induced in normal B6D2F1 mice (reviewed in [2–4]. A single transfer of female parental strain DBA/2 (D2) splenocytes into non-irradiated female B6D2 F1 (D2→F1) hosts results in a chronic graft-vs.-host disease (GVHD) characterized by host B cell expansion, anti-DNA ab production and lupus-like ICGN [5–7]. A single transfer of male D2 splenocytes (m→m), results in a disease that is milder and transient [6]. By contrast, a single transfer of splenocytes from the C57BL/6 (B6) parent (B6→F1) results in an acute GVHD characterized by B6 donor CD8 CTL elimination of host lymphocytes and immunodeficiency lasting months [8]. Sex differences in B6→F1 acute GVHD have not been fully characterized.

Both acute and chronic GVHD are initiated by donor CD4 T cell activation following recognition of allogeneic host MHC II [9–11]. In B6→F1 acute GVHD, B6 donor CD4 T cells produce a strong initial IL-2 response, which in turn promotes a Th1-dominant response consisting of robust TNF and IFNγ production. This is followed by maturation of donor CD8 T cells into short lived effector cell (SLEC) cytotoxic T lymphocytes (CTL) that go on to eliminate F1 splenocytes, particularly B cells [3, 12]. Alloactivated B6 CD4 T cells also provide help to F1 B cells, resulting in B cell expansion over the first week followed by a precipitous decline as they are eliminated by mature B6 CD8 CTLs. In D2→F1 chronic GVHD, D2 mice have a naturally occurring defect in initial IL-2 production that results in skewing of the initial donor CD4 T cell response away from a Th1 program, instead promoting the differentiation of T follicular helper (Tfh) cells and enhanced IgG antibody production [3, 12]. The weak Th1 response nevertheless results in detectable but low level D2 CD8 CTL that are numerically unable to eliminate activated F1 B cells prior to their downregulation [13]. Continued D2 CD4 Tfh cell help to F1 B cells results in unchecked B cell expansion, autoantibody production and eventually lupus-like immune complex glomerulonephritis (ICGN) [3, 6]. The p→F1 model using B6D2 F1 mice thus allows for the comparison of in vivo immune responses in the setting of either a strong Th1-dominated response driven by high IL-2 production (B6 GVHD), or a Tfh-dominated response in the presence of low IL-2 levels (D2 GVHD). This IL-2-dependent bifurcation in CD4 T cell differentiation is consistent with prior studies demonstrating the IL-2-mTORC1 signaling axis as a critical governor of Th1 versus Tfh balance during acute infection, based on differences in metabolic activity [14].

As a result of initial donor CD4 T cell immune skewing, the two forms of GVHD exhibit strikingly divergent phenotypes at two weeks after donor cell transfer, as determined by either gene array [12] or flow cytometric analysis of splenic subsets that can be used as early surrogate markers for longer term phenotypes [13]. Specifically, acute GVHD mice exhibit engraftment of both CD4 and CD8 donor T cells with profound depletion of F1 B cells, whereas chronic GVHD mice exhibit significant engraftment of only the donor CD4 T cell subset coupled with significant expansion of F1 B cells and autoantibody production.

Previous studies have demonstrated that ICGN in D2→F1 mice is more severe in females than males and associated with ∼2-fold greater donor CD4 T cell engraftment at two weeks in females versus males despite transfer of equal numbers of D2 CD4 T cells [5, 6, 7.]. In light of greater emphasis on sex as an important biological variable, affecting immune responses to both self and foreign antigens, we sought to more fully characterize sex differences in both D2→F1 and B6→F1 mice by comparing a sub-threshold and supra-threshold dose of donor cells. By expanding the number of early outcome variables in both the lymphoid and myeloid splenic compartments, we aimed to determine if other heretofore unrecognized sex differences are present not only in the antibody driven immune response in chronic D2 GVHD but also in the generation of a CD8 CTL effector response in acute B6 GVHD mice. Our results indicate that significant sex differences observed in chronic GVHD involve a larger number of lymphoid and myeloid parameters than previously reported.

Significant sex differences in acute GVHD are less extensive, but nevertheless are present. In both forms of GVHD, many of these sex differences have surprisingly large effect sizes, underscoring the significant impact of sexual dimorphism in shaping adaptive immune responses.

## 2. Materials and Methods

### 2.1 Mice

Male and female C57BL6/J (H-2^b^) (B6), DBA/2 (H-2^d^) (D2) and *B6D2F1/J* (H-2^b/d^) (BDF1) mice were purchased from The Jackson Laboratory (Bar Harbor, ME) and housed in our animal facility until testing which was 8 weeks of age. All procedures were pre-approved by the Institutional Animal Care and USE Committee at the Uniformed Services University of Health Sciences.

### 2.2. Induction of GVHD and protocol

GVHD was induced as described [15]. Briefly, single-cell suspensions were prepared in RPMI-1640 medium from the spleens of B6 or D2 donor mice. Cell suspensions were filtered through sterile nylon mesh filters, centrifuged and resuspended in RPMI-1640. The viable cells were counted in a hemocytometer and donor cells adjusted for desired concentration of cells per milliliter. CD4 T cell, CD8 T cell and B220 positive percentages and numbers were quantified using flow cytometry. Disease severity is in the p→F1 model is directly related to the number of donor T cells transferred [16–18]. To be sure our results were in the linear range we tested two doses of unfractionated B6 or D2 donor splenocytes containing either: a) a sub-threshold dose of 4.5 x 10^6^ (4.5M) CD8 T cells; or b) a threshold dose of 5.0 x10^6^ (5M) CD8 T cells [12, 13, 16]. GVHD was induced by tail-vein injection of donor cell suspension normalized to contain either 4.5 x 10^6^ or 5 x 10^6^ CD8 T cells.

Uninjected B6D2F1/J mice that were sex and age matched were used as negative controls. GVHD phenotype was assessed at day 14 by flow cytometric measurement of: a) F1 splenocytes (total versus lymphoid and myeloid subpopulations; and b) donor T cell engraftment. Both sexes were tested using sex-matched and closely age-matched donor and host mice (m→M and f→F). The donor CD4:CD8 ratio was 1.7 for all groups.

### 2.3 Flow cytometry

Spleen cells were first incubated with anti-murine FcγRII/III mAb (clone 2.4G2) for 10 min to block Fc receptors, and then stained with saturating concentrations of Alexa Fluor 647- conjugated, PE-conjugated, FITC-conjugated, PerCPCy5.5-conjugated and Pacific Blue-conjugated mAb against CD4, CD8, B220, H-2K^d^, H-2K^b^, I-A^d^, I-A^b^, CD11c, CD11b, purchased from either BD Biosciences (San Jose, CA), BioLegend (San Diego, CA), eBioscience (San Diego, CA) or Invitrogen (Carlsbad, CA). Cells were fixed in 1% paraformaldehyde prior to analysis. Lymphocytes, monocytes and macrophages were gated by forward and side scatter and fluorescence data collected for at least 10,000 cells. Using previously published gating strategies [13], donor T cell populations were identified as CD4 and CD8 -/+ for MHC class I of the uninjected parent (H2-K^d^ or H2-K^b^ negative). Donor and host CD4 and CD8 T cells were further analyzed for upregulation of CD44 which exhibited a typical bimodal distribution denoting CD44^hi^ or CD44^low^ populations. Donor and host CD4 Tfh cells were assessed as CD3+, CD4+, ICOS+, CXCR5+ based on previously published gating strategy [19]. Host B cells were gated as B220+ and MHC class II+ of the uninjected parent (I-A^d^ or I-A^b^ positive). Host dendritic cells (DCs) and macrophages were identified as MHC class II+ and CD11c+ or CD11b+, respectively, using a broad gate and previously published gating strategies [7, 12, 19, 20]. Following completion of the staining protocol, cells were analyzed immediately using a BD LSRII flow cytometer (BD Biosciences, San Jose, CA).

### 2.4 Statistical Analysis

Statistical comparisons were performed using Prism 9.4 (GraphPad Software, San Diego, CA 92108). Mice were tested individually, and data are expressed as group mean ± SEM. Statistical significance comparing two groups was determined using Student’s t-test. p values <0.05 were considered statistically significant. Effect sizes are reported as Cohen’s d, calculated as the difference in means divided by the pooled standard deviation. In this context, we considered effect sizes > 1.0 to be large effects.

## 3. Results

### 3.1 B6→F1 acute GVHD: splenocyte subset changes

#### 3.1.1 Dose effects in males

For m→M mice, the 5M CD8 T cell inoculum resulted in large and significant reductions of recipient cell populations relative to uninjected control F1 mice for the following host F1 parameters: total splenocytes, B cells, CD4 T cells, CD8 T cells and CD11b cells (Fig 1A-1E, Table 1A). Similarly, the 4.5M cell dose also resulted in statistically significant reductions vs. control F1 mice for these same parameters, except for host CD8 T cells (Fig 1A-1E, Table 1A). As expected, F1 B cells exhibited the most profound reductions and the highest effect sizes (d>40) at both the 5M dose (0.7% of control) and 4.5M dose (2.0% of control) (Table 1A). Reductions for the other subsets were less severe and ranged from 29% - 90% of control (Table 1A). At the 5M dose, there was a hierarchy of host splenocyte depletion severity with B cells being the most severely depleted vs. control (0.7%), followed by CD11b+ macrophages (29.6%) and then CD4 (50.7%) and CD8 T cells (67.8%). A similar hierarchy was observed at the 4.5M dose, with less severe depletion of CD11b+ cells. In contrast, host CD11c+ DC numbers behaved differently from the other F1 splenocyte subsets; we observed no significant reductions in this population compared to controls, and values for the 4.5M dose were significantly increased versus either control or the 5M dose.

**Figure 1.**
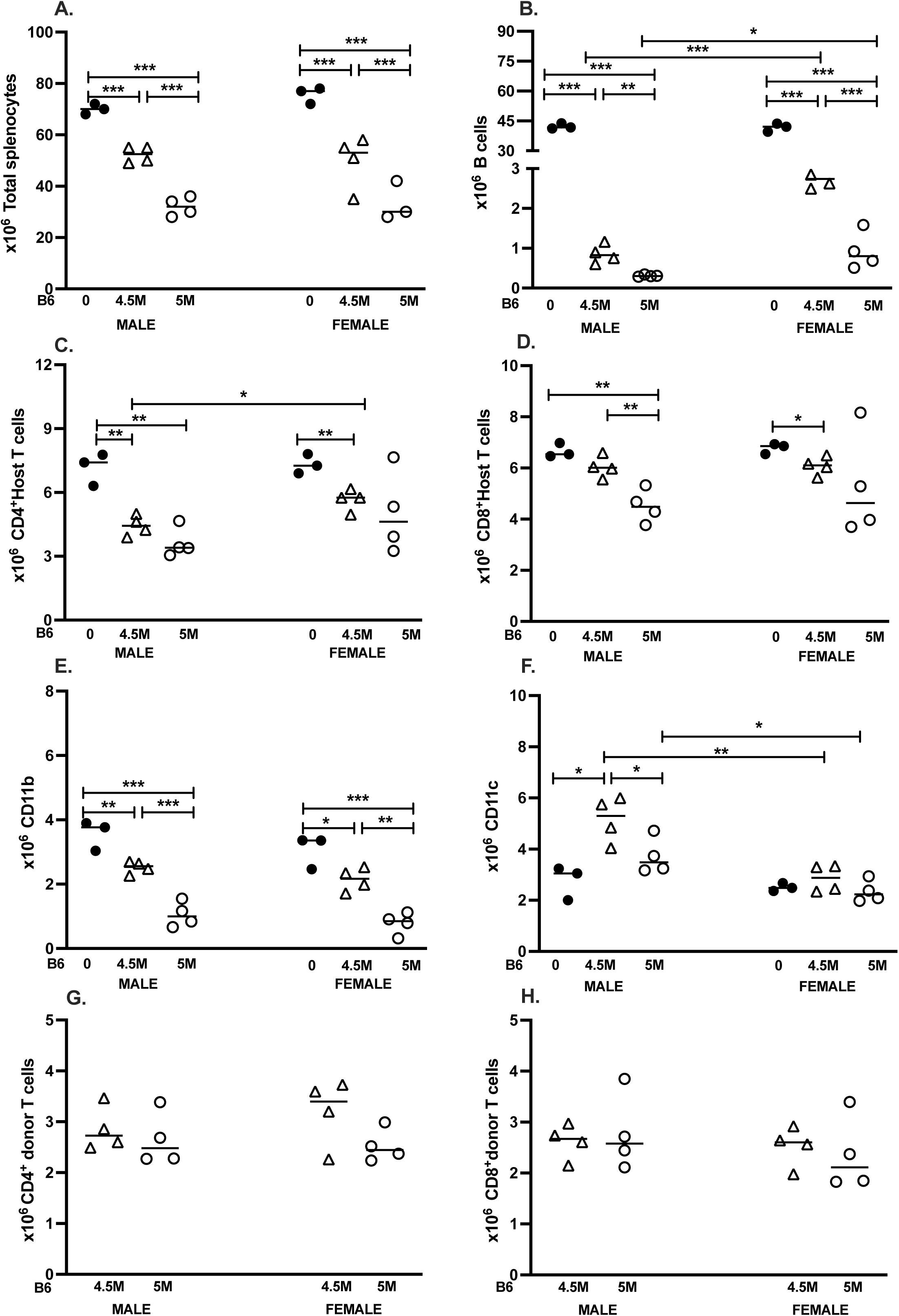
Dose response and sex differences for B6→F1 GVHD phenotype at two weeks. BDF1 mice were either uninjected (dark circles) or received unfractionated splenocytes from B6 donors normalized to contain either 4.5 x 10^6^ CD8 T cells (4.5M)(open triangles) or 5 x 10^6^ CD8 T cells (5M) (open circles) as described in Methods. Donors and hosts were either both males or both females. Spleens were harvested at two weeks after donor cell transfer, and splenocytes were assessed by flow cytometry to quantify: (A) total splenocytes; (B) host B cells; (C) host CD4 T cells; (D) host CD8 T cells; (E) host CD11b+ macrophages; (F) host CD11c+ DCs; (G) donor CD4 and (H) donor CD8 T cell engraftment. Values represent group mean ± SE (n= 4/grp experimental; n=3/grp controls). For all figures, *p<0.05, **p<0.01, ***p<0.001. Male and female donor splenocyte inoculum contained either 7.7 or 8.6 x 10^6^ CD4 T cells for the 4.5M and 5M doses respectively.

**Table 1.**
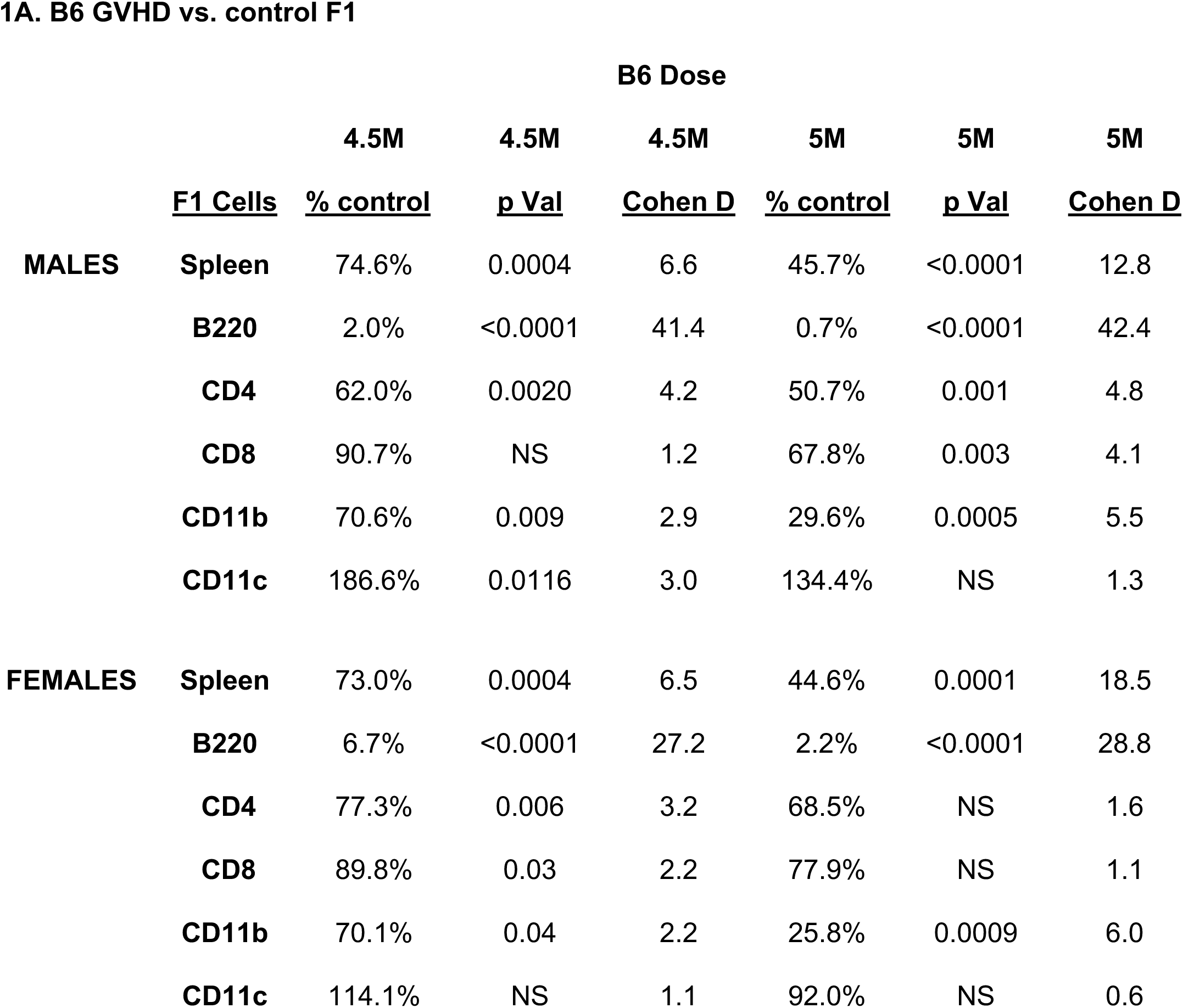
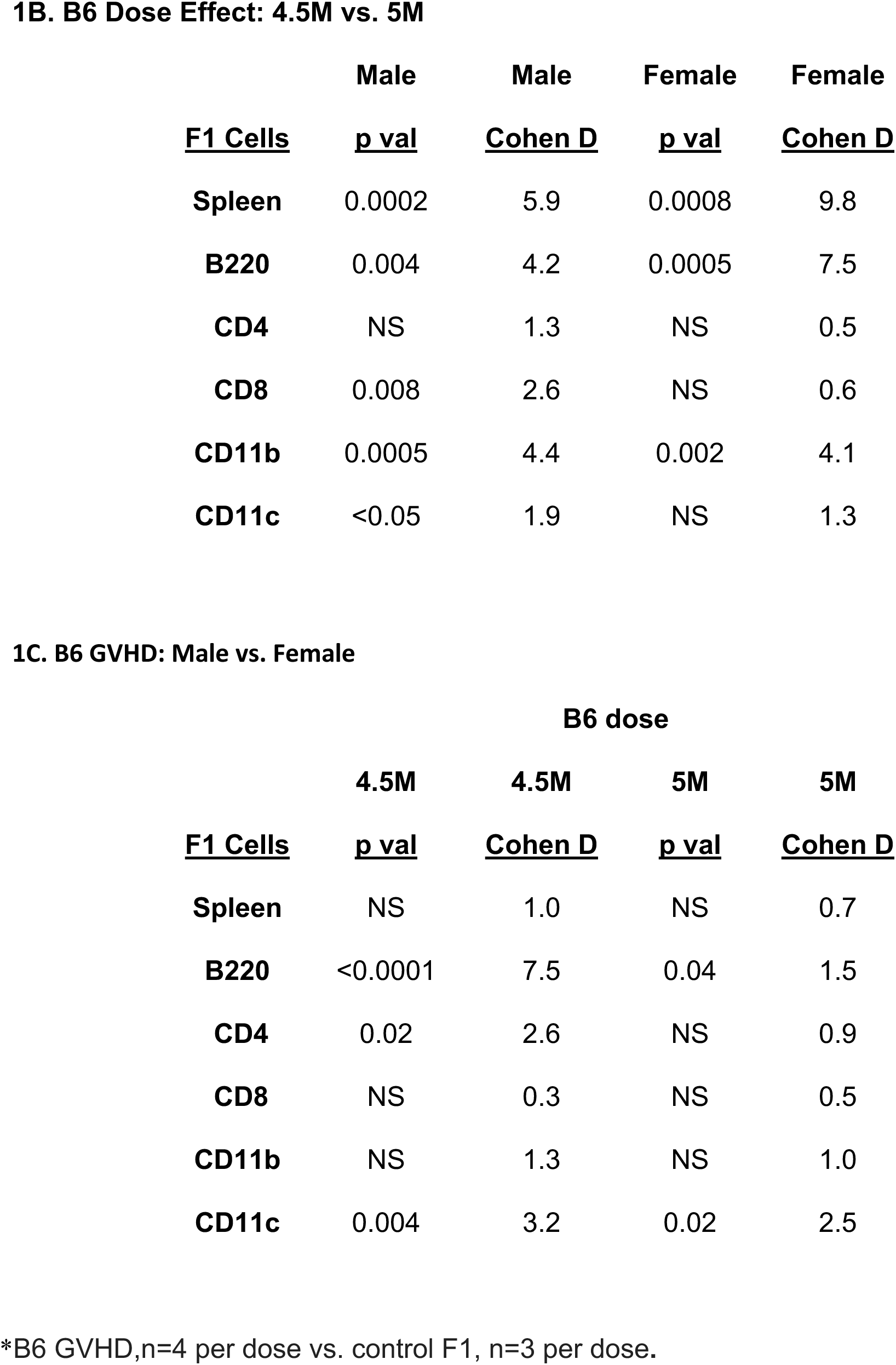
Statistical Comparisons for Acute B6 GVH*.

**1A. B6 GVHD vs. control F1**
|  |  | **B6 Dose** |  |  |  |  |  |
| --- | --- | --- | --- | --- | --- | --- | --- |
|  |  | **4.5M** | **4.5M** | **4.5M** | **5M** | **5M** | **5M** |
|  | **<u>F1 Cells</u>** | **<u>% control</u>** | **<u>p Val</u>** | **<u>Cohen D</u>** | **<u>% control</u>** | **<u>p Val</u>** | **<u>Cohen D</u>** |
| **MALES** | **Spleen** | 74.6% | 0.0004 | 6.6 | 45.7% | <0.0001 | 12.8 |
|  | **B220** | 2.0% | <0.0001 | 41.4 | 0.7% | <0.0001 | 42.4 |
|  | **CD4** | 62.0% | 0.0020 | 4.2 | 50.7% | 0.001 | 4.8 |
|  | **CD8** | 90.7% | NS | 1.2 | 67.8% | 0.003 | 4.1 |
|  | **CD11b** | 70.6% | 0.009 | 2.9 | 29.6% | 0.0005 | 5.5 |
|  | **CD11c** | 186.6% | 0.0116 | 3.0 | 134.4% | NS | 1.3 |
| **FEMALES** | **Spleen** | 73.0% | 0.0004 | 6.5 | 44.6% | 0.0001 | 18.5 |
|  | **B220** | 6.7% | <0.0001 | 27.2 | 2.2% | <0.0001 | 28.8 |
|  | **CD4** | 77.3% | 0.006 | 3.2 | 68.5% | NS | 1.6 |
|  | **CD8** | 89.8% | 0.03 | 2.2 | 77.9% | NS | 1.1 |
|  | **CD11b** | 70.1% | 0.04 | 2.2 | 25.8% | 0.0009 | 6.0 |
|  | **CD11c** | 114.1% | NS | 1.1 | 92.0% | NS | 0.6 |

1B. B6 Dose Effect: 4.5M vs. 5M
|  | Male | Male | Female | Female |
| --- | --- | --- | --- | --- |
| <u>F1 Cells</u> | <u>p val</u> | <u>Cohen D</u> | <u>p val</u> | <u>Cohen D</u> |
| Spleen | 0.0002 | 5.9 | 0.0008 | 9.8 |
| B220 | 0.004 | 4.2 | 0.0005 | 7.5 |
| CD4 | NS | 1.3 | NS | 0.5 |
| CD8 | 0.008 | 2.6 | NS | 0.6 |
| CD11b | 0.0005 | 4.4 | 0.002 | 4.1 |
| CD11c | <0.05 | 1.9 | NS | 1.3 |

1C. B6 GVHD: Male vs. Female
|  | B6 dose |  |  |  |
| --- | --- | --- | --- | --- |
|  | 4.5M | 4.5M | 5M | 5M |
| <u>F1 Cells</u> | <u>p val</u> | <u>Cohen D</u> | <u>p val</u> | <u>Cohen D</u> |
| Spleen | NS | 1.0 | NS | 0.7 |
| B220 | <0.0001 | 7.5 | 0.04 | 1.5 |
| CD4 | 0.02 | 2.6 | NS | 0.9 |
| CD8 | NS | 0.3 | NS | 0.5 |
| CD11b | NS | 1.3 | NS | 1.0 |
| CD11c | 0.004 | 3.2 | 0.02 | 2.5 |

\*B6 GVHD, n=4 per dose vs. control F1, n=3 per dose.

Comparing the two donor cell doses to each other, significantly greater reduction of host cell subsets was noted for the 5M dose for all parameters in Fig. 1A-1E except for F1 CD4 T cells, where the GVHD-associated reductions did not differ significantly between the two doses (Fig. 1C, Table 1B). For F1 B cells, the 5M and 4.5M dose resulted in maximal (>99%) and near maximal (98%) depletion, respectively, whereas sub-maximal depletion was seen for all other splenic subsets.

#### 3.1.2 Dose effects in females

For f→F mice, all host splenocyte subsets in Fig. 1A-1E were significantly reduced vs. control F1 mice, apart from F1 CD4 and CD8 T cells (5M dose only) where decreases did not reach significance (Table 1A). Subset ranking according to the percent reduction vs. control yielded the same aforementioned hierarchy for males, with female F1 B cells showing the most profound reductions at both the 5M (2.2% of control) and 4.5M dose (6.7% of control) and very large effect sizes (d >27, Table 1), followed by F1 CD11b+ macrophages (25.8% of control), F1 CD4 and CD8 T cells (68.5% and 77.9% of control respectively) (Table 1A). As with male acute GVHD, host CD11c+ DC numbers in f→F mice behaved differently from the other splenic subsets and exhibited no significant change from control F1 levels at either dose (Fig. 1F).

Comparing the two donor cell doses to each other in f→F mice, the 5M dose resulted in significantly greater reductions for total splenocytes, F1 B cells and F1 CD11b+ macrophages with the largest effect sizes seen for total splenocytes and F1 B cells (Table 1B). A trend toward greater reduction in F1 CD4 and CD8 T cells was noted for the 5M vs. 4.5M dose, but did not reach statistical significance.

#### 3.1.3. **Sex differences**

Control uninjected F1 values for all 6 parameters did not differ between the sexes (Fig. 1A-1F). Compared to f→F mice, m→M mice exhibited significantly greater reductions in host B cells (both doses) and host CD4 T cells (4.5M dose only) (Table 1C). No sex-based differences at either dose were seen for total splenocytes, host CD8 or host CD11b+ macrophages. Interestingly, F1 CD11c+ DC cells were significantly increased in males vs. females at both doses. There were no significant dose or sex related differences in the engraftment of either donor CD4 or donor CD8 T cells (Fig. 1G, 1H).

#### 3.1.4 **Summary.**

Overall, both doses of donor CD8 T cells from both sexes resulted in significant differences vs. control F1, with very large effect sizes. We detected a hierarchy of splenic subset depletion, with F1 B cells consistently the most sensitive to donor T cell elimination and exhibiting the greatest percentage reduction and largest effect size ( >41.0 males; >27.0 females at both doses, Table 1A. There were no sex differences in the hierarchy of F1 splenic subset depletion. The significant differences observed between the two donor CD8 T cell doses demonstrate that even a mild reduction in CD8 T cell effectors translates to an increase in host splenic subpopulations, with F1 B cells being the most sensitive indicator.

### 3.2 D2→F1 chronic GVHD: splenocyte subset changes

#### 3.2.1 Dose effects in males

For m→M mice, both the 5M and 4.5M donor CD8 T cell doses resulted in an increase in recipient cell populations vs. control F1 mice for the following host F1 parameters: total splenocytes, B cells, CD4 T cells, CD8 T cells, CD11b and CD11c cells (Fig 2A-2F, Table 2A). These increases were significant for all groups except: a) F1 CD11c+ at 4.5 M dose; and b) F1 CD8 T cell values, which were not significantly different from F1 values at the 5M dose, and surprisingly were significantly reduced at the 4.5M dose. Unlike the acute GVHD model in which F1 B cells are most profoundly affected, in D2→F1 mice the percentage increase over control was highest at both doses for F1 CD4 T cells (>200%), followed in descending order by F1 CD11c+ DCs, CD11b+ macrophages, B cells and CD8 T cells (Table 2A). Large effect sizes were measured for total splenocytes and for F1 CD4 T cells at both 4.5M (d >7.0) and 5M (d >11.0) doses (Table 2A). F1 CD8 T cells were the exception, showing a decrease at the 4.5M dose and only a 118% increase at the 5M dose vs. control.

**Figure 2.**
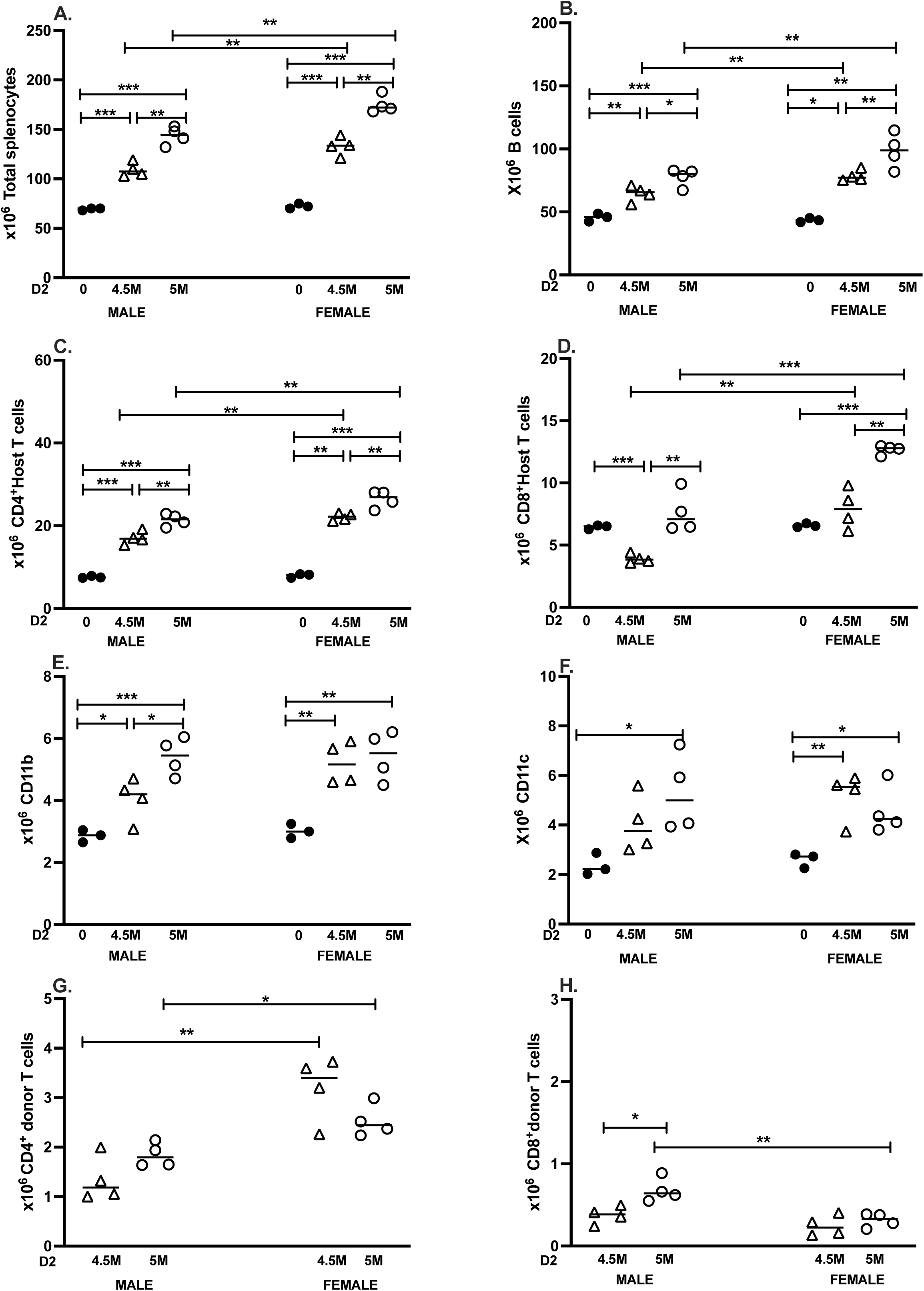
Dose response and sex differences for D2→F1 GVHD phenotype at two weeks. BDF1 mice were either uninjected (dark circles) or received unfractionated splenocytes from D2 donors normalized to contain either 5 x 10^6^ CD8 T cells (open circles) or 4.5 x 10^6^ CD8 T cells (open triangles) as described in Methods. Donors and hosts were either both either males or both females. Spleens were harvested at two weeks after donor transfer, and splenocytes were assessed by flow cytometry to quantify: (A) total splenocytes; (B) host B cells; (C) host CD4 T cells; (D) host CD8 T cells; (E) host CD11b cells; (F) host CD11c cells; (G) donor CD4 and (H) donor CD8 T cell engraftment. Values represent group mean ± SE (n= 4/grp experimental; n=3/grp controls). For all figures, *p<0.05, **p<0.01, ***p<0.001. For the 4.5M and 5M doses, the male splenocyte donor inoculum contained 8.2 or 9.2 x 10^6^ CD4 T cells respectively, and females received either 7.8 or 8.6 x 10^6^ CD4 T cells respectively.

**Table 2.** Statistical Comparisons for Chronic D2 GVH*.

**2A. DBA GVHD vs. control F1**
|  |  | **<u>DBA dose</u>** |  |  |  |  |  |
| --- | --- | --- | --- | --- | --- | --- | --- |
|  |  | **4.5M** | **4.5M** | **4.5M** | **5M** | **5M** | **5M** |
|  | **<u>F1 Cells</u>** | **<u>% control</u>** | **<u>p VALUE</u>** | **<u>Cohen D</u>** | **<u>% control</u>** | **<u>p VALUE</u>** | **<u>Cohen D</u>** |
| **MALES** | **Spleen** | 157.6% | 0.0002 | 7.9 | 207.0% | <0.0001 | 11.4 |
|  | **B220** | 141.4% | 0.0054 | 3.8 | 170.1% | 0.0008 | 5.7 |
|  | **CD4** | 224.6% | 0.0002 | 8.2 | 281.2% | <0.0001 | 12.7 |
|  | **CD8** | 60.6% | <0.0001 | 8.2 | 118.1% | NS | 0.9 |
|  | **CD11b** | 141.6% | 0.04 | 2.1 | 189.5% | 0.001 | 5.6 |
|  | **CD11c** | 169.9% | NS | 1.8 | 223.0% | 0.03 | 2.4 |
| **FEMALES** | **Spleen** | 183.7% | 0.0001 | 8.8 | 242.3% | <0.0001 | 15.7 |
|  | **B220** | 183.9% | 0.0416 | 10.3 | 239.0% | 0.0093 | 8.4 |
|  | **CD4** | 278.2% | 0.0015 | 19.5 | 330.0% | <0.0001 | 12.1 |
|  | **CD8** | 120.7% | NS | 1.1 | 192.4% | <0.0001 | 19.3 |
|  | **CD11b** | 172.9% | 0.0032 | 4.2 | 180.0% | 0.0041 | 4.1 |
|  | **CD11c** | 199.3% | 0.0075 | 3.5 | 176.9% | 0.0216 | 2.7 |

2B. DBA Dose Effect: 4.5M vs. 5M
|  | Male | Male | Female | Female |
| --- | --- | --- | --- | --- |
| <u>F1 Cells</u> | <u>p VALUE</u> | <u>Cohen D</u> | <u>p VALUE</u> | <u>Cohen D</u> |
| **Spleen** | 0.001 | 4.2 | 0.0006 | 4.6 |
| **B220** | 0.0341 | 1.9 | 0.0059 | 3.1 |
| **CD4** | 0.0078 | 2.8 | 0.0095 | 2.6 |
| **CD8** | 0.0047 | 3.0 | 0.0012 | 4.1 |
| **CD11b** | 0.0249 | 2.1 | NS | 0.2 |
| **CD11c** | NS | 0.9 | NS | 0.7 |

2C. DBA GVHD: Male vs. Female
|  | DBA dose |  |  |  |
| --- | --- | --- | --- | --- |
|  | 4.5M | 4.5M | 5M | 5M |
| <u>F1 Cells</u> | <u>p VALUE</u> | <u>Cohen D</u> | <u>p VALUE</u> | <u>Cohen D</u> |
| **Spleen** | 0.007 | 2.8 | 0.003 | 3.5 |
| **B220** | 0.01 | 2.7 | 0.009 | 3.0 |
| **CD4** | 0.002 | 3.9 | 0.008 | 2.7 |
| **CD8** | 0.003 | 3.4 | 0.001 | 4.1 |
| **CD11b** | NS | 1.4 | NS | 0.0 |
| **CD11c** | NS | 1.2 | NS | 0.5 |

\*DBA GVHD, n=4 per dose vs. control F1, n=3 per dose.

Comparing the two donor cell doses to each other, the 5M dose resulted in a greater increase in all F1 parameters vs. 4.5M dose (Fig. 2A-2F), and was statistically significant for all except for host CD11c+ DCs and CD11b+ macrophages at the 5M dose (Table 2B).

#### 3.2.2. Dose effects in females

For f→F mice, both donor cell doses resulted in a statistically significant increase in all host splenocyte subsets (Fig. 2A-2F) except host CD8 T cells at the 4.5M dose. Very large effect sizes (d=8-19) were seen for many of the parameters (Table 2A). F1 CD8 T cells exhibited the lowest percentage increase (120.7%) over control vs. F1 at the 4.5M dose. However, at the 5M dose, the percentage increase (192.4%) was comparable to F1 CD11b+ and CD11c+ cells. As with male DBA GVHD mice, F1 CD4 T cells exhibited the highest percentage increase vs. control at either dose. At the 5M dose, F1 B cells were the second highest and F1 CD11c, CD11b and CD8 T cell were <200% increased.

Comparing the two donor cell doses to each other, the 5M dose resulted in significantly greater expansion of F1 total splenocytes, B cells, CD4 and CD8 T cells, with the percentage increase over F1 control ranging from 192% to 330% (Table 2B). Effect sizes ranged from d=2-4 at both doses. Expansion was also seen for host CD11b+ macrophages and CD11c+ DCs, but did not differ significantly between the two doses (Fig. 2E, 2F, Table 2B).

#### 3.2.3. Sex differences

Control F1 values for all 6 parameters did not differ between the sexes (Fig. 2A-2F). Female D2 GVHD mice exhibited significantly greater numbers of F1 splenocytes, B cells, CD4 T cells and CD8 T cells at either donor cell dose compared to m→M mice, (Fig. 2A-2D) with effect sizes in the 2-4 range (Table 2C). There were no significant sex differences seen for either the CD11b+ or CD11c+ subsets.

#### 3.2.4 Donor T cell engraftment

##### CD4 T cells

There were no significant dose-related differences in donor D2 CD4 engraftment in same-sex comparisons; however, f→F1 mice exhibited significantly greater donor CD4 engraftment vs. m→M mice at both doses (Fig. 2G), consistent with previous reports [5, 7]. Interestingly, the effect size was greater at the 4.5M dose (d=3.7) vs. the 5M dose (d=2.4). <u>CD8 T cells.</u> Significant differences in D2 donor CD8 engraftment were observed only at the 5M dose, where both dose effects and sex differences were noted (Fig. 2H). CD8 engraftment for m→M mice was significantly greater at the 5M dose vs. the 4.5M dose in males, and compared to either dose in females. As previously reported [13], total donor CD8 T cell engraftment is reduced for both males and females (≤ 1 x 10^6^) compared to that of B6→F1 (≥2 x 10^6^) mice due to a failure of D2 CD8 T cell expansion (Fig. 2H).

Taken together, data summarized in Figures 1 and 2 demonstrate that transfer of an unfractionated donor splenocyte inoculum containing 5 x 10^6^ CD8 T cells is sufficient to induce near maximal reductions in host B cell populations for B6→F1 acute GVHD and >200% increase in several splenic subsets in D2→F1 chronic GVHD mice. Moreover, attenuation of host cell phenotypes in either model is readily detectable at a dose of 4.5M donor CD8 T cells, with significant differences noted in several splenocyte subpopulations. Although F1 B cells are the most sensitive to elimination in B6 GVHD mice, they rank second to F1 CD4 T cells in D2 GVHD subset expansion.

### 3.3. Upregulation of CD44 on T cells in the B6 GVHD model

To gauge T cell activation in both GVHD models, we also measured the percentage and absolute number of CD44^hi^ donor and host CD4 and CD8 T cells (Figs 3A-3H).

**Figure 3.**
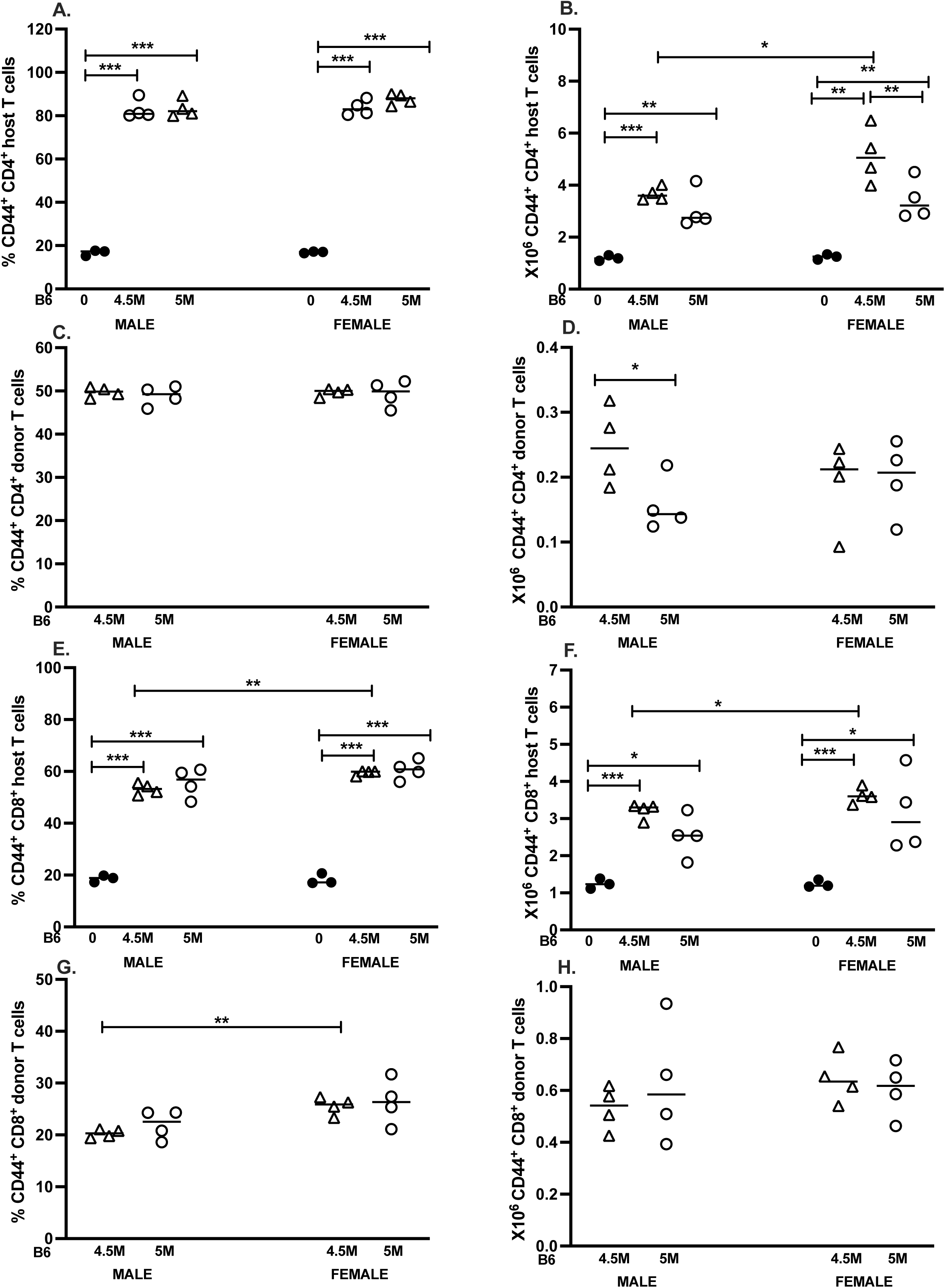
T cell CD44 upregulation in B6 GVHD mice is similar in both males and females. The cohort described in Figure 1 was analyzed at day 14 for CD44 expression on: 1) F1 and donor (B6) CD4 T cells from control F1 (A, B), and B6 GVH mice (C,D) and 2) F1 and donor (B6) CD8 T cells from control F1 (E,F) and B6 GVH mice (G,H). Results are shown as group mean ± SE for the percentage (A,C,E.G) or total number (B,D, F, H) of CD44^hi^ cells.

#### F1 CD4 T cells

The percentage of F1 CD44^hi^ CD4 T cells was significantly greater than controls for both donor cell doses and both sexes (∼80%), and did not differ significantly between the two doses or between sexes (Fig. 3A). Similarly, the numbers of F1 CD44^hi^ CD4 T cells were also greater than control F1 values for both GVHD doses and for both sexes (Fig. 3B). Sex differences were noted only at the 4.5M dose, where absolute numbers of CD44^hi^ F1 CD4 T cells were significantly greater for females vs. males.

#### Donor B6 CD4 T cells

The percentage of CD44^hi^ B6 donor CD4 T cells (∼50%) did not differ significantly between doses or sexes (Fig. 3C). Males exhibited significantly greater numbers of B6 donor CD44^hi^ CD4 T cells at the 4.5M dose vs. 5M dose. There were no significant sex differences in numbers of B6 CD44hi CD4 T cells.

#### F1 CD8 T cells

Both males and females exhibited a significant increase in the percent and numbers of F1 CD44^hi^ CD8 T cells vs. control F1 mice (Fig. 3F, 3G). Intriguingly, a small but significant increase (p=0.04) in the percentage and total number of CD44^hi^ F1 CD8 T cells was observed in females at the 4.5M inoculum.

#### B6 donor CD8 T cells

There were no significant differences between same-sex comparisons of the percentage of CD44^hi^ donor CD4 T cells. However, a slightly higher percentage of CD44^hi^ donor CD4 T cells was measured in females for the 4.5M dose (Fig. 3G), although no significant differences in absolute cell number were detected for either dose or between sexes (Fig. 3H).

Overall, robust activation was observed for donor and host CD4 and CD8 T cells as measured by CD44 upregulation at two weeks post-transfer, and was generally similar in both males and females. Of note, significant sex differences were discernible at the 4.5M dose, including: a) numbers of CD44^hi^ F1 CD4 T cells; b) percent and number of F1 CD44^hi^ CD8 T cells; and c) the percent of donor CD8 T cells. The absence of dissimilarity at the maximal 5M dose suggests that sex differences in acute GVHD are subtle and best seen at sub-maximal doses of donor cells. Activation of host CD8 T cells is consistent with a host-vs.- graft (HVG) response as previously described [13].

### 3.4 Upregulation of CD44 on T cells in the D2 GVHD model

#### F1 CD4 T cells

F1 CD44^hi^ CD4 T cells were expanded (both % and number) in D2 GVHD mice, and values were significantly greater than control F1 at both doses (5M > 4.5M) (Fig. 4A, 4B). Additionally, females exhibited significantly greater numbers of CD44^hi^ F1 CD4 T cells vs. males at both doses. Thus, F1 CD4 T cells exhibit not only a significant increase in absolute cell numbers (Fig. 2C), but also in activation status.

**Figure 4.**
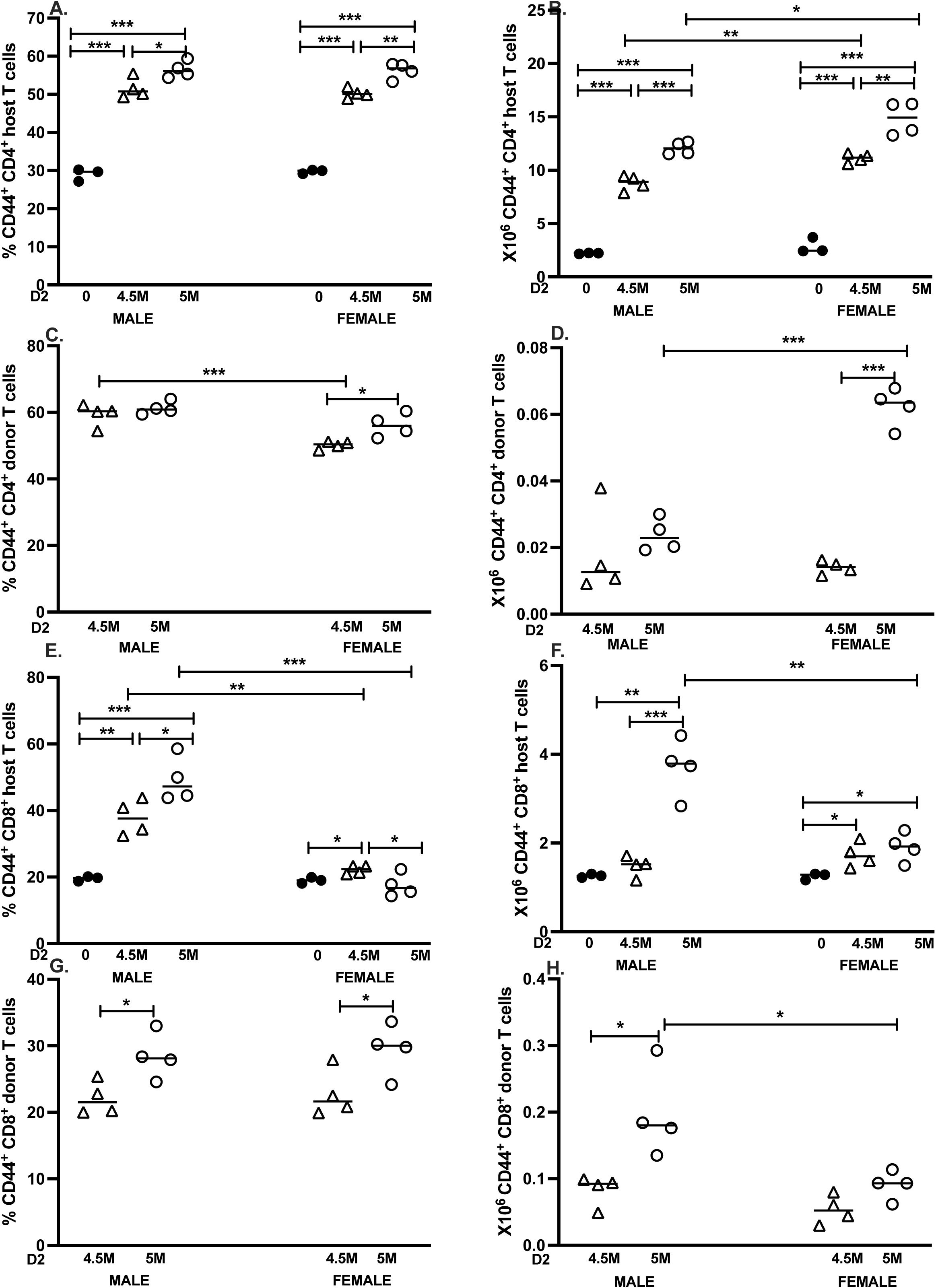
T cell CD44 upregulation in D2 GVHD mice: sex differences are seen in donor and host T cell CD44 upregulation. The cohort described in Figure 2 was analyzed at day 14 for CD44 expression on: 1) F1 and donor (D2) CD4 T cells from control F1 (A, B), and D2 GVH mice (C,D) and 2) F1 and donor (D2) CD8 T cells from control F1 (E,F) and D2 GVH mice (G,H). Results are shown as group mean ± SE for the percentage (A,C,E.G) or total number (B,D, F, H) of CD44^hi^ cells.

#### D2 Donor CD4 T cells

The percentage of CD44^hi^ donor CD4 cells was comparable in males at both doses; however, the percentage was significantly lower at the 4.5M dose vs. 5M in females (Fig. 4C). At the 4.5M dose, male percentages were significantly greater than females. Nevertheless, females exhibited greater absolute numbers of CD44^hi^ donor CD4 T cells at the 5M dose (Fig.4D).

#### F1 CD8 T cells

Males demonstrated a significant increase in the percentage of F1 CD44^hi^CD8 T cells at both doses vs. control (5M > 4.5M) (Fig. 4E). By contrast, females exhibited a much smaller yet significant change in the percentage of CD44^hi^ F1 CD8 T cells vs. control at the 4.5M dose only, with no significant difference vs. control at the 5M dose. Hence significant sex differences were found for F1 CD44^hi^ CD8 T cells, with males exhibiting a significantly greater percentage of CD44^hi^ F1 CD8 T cells vs. females at both doses (Fig. 4E). Similarly, the absolute number of male CD44^hi^ F1 CD8 T cells were significantly elevated vs. control and vs. females only at the 5M dose, whereas females showed much smaller yet significant elevations at both doses (Fig. 4F).

#### D2 Donor CD8 T cells

Both males and females exhibited significantly greater percentages of CD44^hi^ donor CD8 T cells at the 5M vs. 4.5M dose, with comparable values recorded between the sexes (Fig. 4G). However, only males demonstrated a significant increase in the absolute number of donor CD44^hi^ CD8 T cells between the two doses, and were significantly elevated compared to females (Fig. 4H).

Together, these data support previous work indicating that in D2 GVHD mice, females exhibit a greater expansion of donor D2 CD4 T cells, whereas males exhibit a stronger CD8 T cell-mediated donor anti-host and reciprocal host anti-donor response [7].

### 3.5 Quantification of CD4 Tfh cells in GVHD mice

#### 3.5.1 CD4 Tfh cells in B6 GVHD mice

We also assessed changes in donor and F1 CD4+ICOS+CXCR5+ Tfh populations in both GVHD models by flow cytometry. In the B6 GVHD model, a significant increase in both the percentage and number of Tfh cells was detected for F1 CD4 T cells vs. control F1 mice at both doses for males and females (Fig. 5A-B). Sex differences were seen only for the numbers of F1 Tfh cells at the 4.5M dose, where F1 females exhibited a significant increase vs. males (Fig 5B). Both males and females also showed a significant dose-dependent increase in the percentage (but not absolute number) of donor Tfh cells, with no significant sex differences observed.

**Figure 5.**
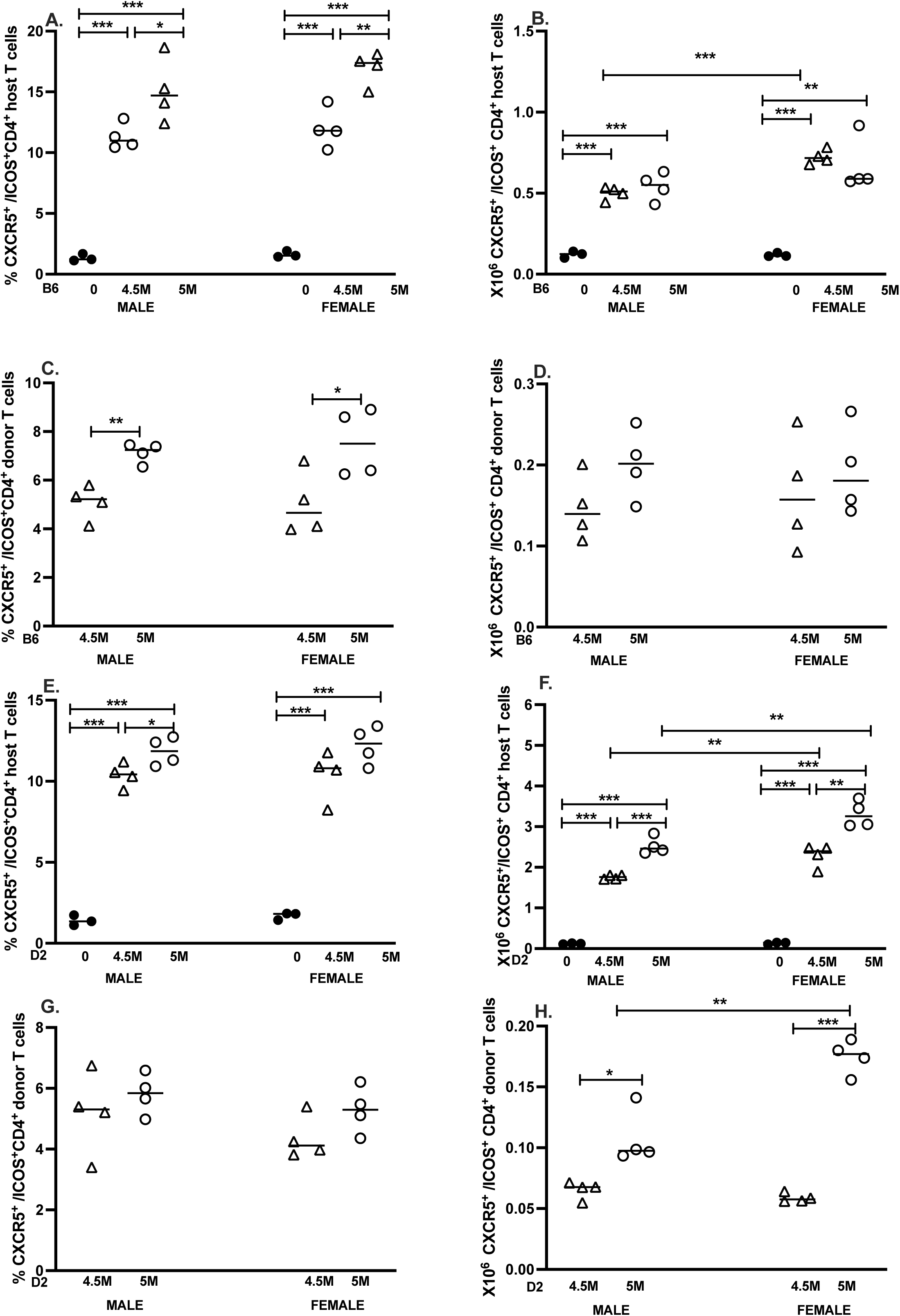
Sex differences in donor and F1 CD4 Tfh cells are seen only in D2 GVHD. The cohorts described in Figures 1 and 2 were further analyzed on day 14 to quantify CD4+ CXCR5+ICOS1+ TFh cells from B6 GVHD mice (A-D) or D2 GVHD mice (E-H). Results for uninjected F1 controls (A,B, E, F) and GVHD mice (C,D, G, H) are shown as group mean ± SE for the percent (A,C,E.G) or total number (B,D, F, H) of CXCR5,ICOS1+ TFh cells.

#### 3.5.2 CD4 Tfh cells in D2 GVHD mice

In the D2 GVHD model, significant increases in the percentage and number of F1 CD4 Tfh cells were noted at both doses in males and females vs. control F1 mice (Fig 5E-F). Additionally, females exhibited significantly greater numbers of F1 CD4 Tfh cells relative to males at both donor cell doses. Conversely, there were no significant differences in the percentage of donor CD4 Tfh cells for same-sex or between-sex comparisons (Fig. 5G). However, both sexes exhibited a significant dose-related increase in the absolute number of donor CD4 Tfh cells, which was significantly greater in females for the 5M dose (Fig. 5H).

In summation, we detected sex differences in B6 GVHD Tfh cells only at the 4.5M cell dose, for which females had significantly greater numbers of F1 CD4 Tfh cells. By contrast, female D2 GVHD mice exhibited greater numbers of F1 CD4 Tfh cells (both doses) and donor D2 CD4 Tfh cells (5M dose only).

## 4. Discussion

To our knowledge, our study is the first to compare sex differences in cellular composition and phenotype for acute and chronic GVHD mice at 2 weeks post-induction, using two different doses of donor splenocytes standardized to CD8 T cell numbers. Our results in acute B6 GVHD mice demonstrate that F1 lymphoid and myeloid splenocyte subpopulations exhibit differential sensitivity to donor CD8 CTL elimination, with F1 B cells being the most dramatically affected. In both males and females, F1 B cells exhibited the greatest percentage reduction (<7% of control) at either donor cell dose compared to other splenic subsets, and were associated with very large effect sizes. Despite near complete elimination of F1 B cells, other F1 splenocyte subsets showed progressively increasing survival in the following rank order at the 5M dose: F1 CD11b+ macrophages (∼26% of control), F1 CD4 T cells (∼69% of control) and F1 CD8 T cells (∼78% of control). Total splenocytes (∼45% control) in B6 GVHD mice reflect the overall combined differential sensitivity to CTL-mediated elimination, with very large effect sizes observed in both males (12.8) and females (18). Long term, splenic repopulation by donor cells proceeds in a different order, with myeloid cells (Mac-1^+^) being the first to appear followed by B cells, CD8 T cells, and CD4 T cells at much later timepoints [8].

The greater sensitivity of B cells to CTL-dependent elimination is underscored in the dose-response comparison (Table 1B), where a 10% decrease in the number of donor CD8 T cells transferred results in a significant increase in surviving F1 B cells, associated with large effect sizes in both males (4.2) and females (7.5). Total F1 splenocyte numbers also exhibited a significant dose-response effect with effect sizes of 5.9 (males) and 9.8 (females). Lastly, CD11b+ macrophages exhibited sub-maximal elimination (∼25%-30% of control) at the 5M dose, with a significant dose response and effect size (∼4) found for both males and females. The remaining subsets exhibited more variability, with F1 CD8 T cells exhibiting a significant dose-dependent effect for males only. No significant dose effect was observed for male or female F1 CD4 subsets, although we recorded a large effect size for males (1.3) and a medium effect size for females (F1 CD4= 0.5; F1 CD8= 0.6). Overall, the proportion of F1 B cells remained the most sensitive quantitative indicator of the presence donor CTL in both males and females. These results are consistent with our previous work showing that F1 B cell numbers are a more sensitive marker for the strength of the donor anti-F1 CD8 CTL response when compared to either in vitro or ex-vivo ^51^Cr release cytotoxic assays [13, 21], with the in vivo CTL assay remaining the most sensitive [13],

Although there were no sex differences in the rank order of subset susceptibility to CTL elimination, significant sex differences were seen in the severity of elimination for F1 B cells and F1 CD4 T cells. Depletion of F1 B cells in females, while also severe (<7% of control), was nevertheless significantly less than that of males at both donor cell doses, accompanied by a large effect size at the 4.5M dose (7.5) and a lesser but significant effect size (1.5) at the 5M dose. Sex differences were also discovered in the elimination of F1 CD4 T cells, but were less robust than those noted for F1 B cells, and seen primarily at the 4.5M dose in which females exhibited significantly less elimination (greater survivability) than males. This trend was associated with greater numbers of surviving female F1 CD44^hi^ and Tfh CD4 T cells vs. males.

There were no significant sex differences in total numbers F1 CD8 T cells, although females exhibited a small but significantly greater number of CD44^hi^ F1 CD8 T cells at the 4.5M dose. Donor anti-F1 CTL GVH reactions induce a counter regulatory host-vs.-graft (HVG) reaction initially termed “hybrid resistance” [22], mediated by both NK cells [23] and F1 CD8 T cells [24, 25]. Indeed, F1 anti-parent CTL have been shown in vitro and in vivo [13, 26, 27]. Using the absolute number of CD44^hi^ F1 CD8 T cells to approximate the strength of CTL-mediated HVG, relative sparing of F1 CD8 T cells in females is consistent with a weaker HVG elicited in response to the weaker donor anti-host CD8 GVH response in females (as measured by reduced F1 B cell elimination vs. males).

By contrast, females exhibited a significantly greater reduction in F1 CD11c+ DCs vs. males at both donor cell doses. Kinetic studies in male acute GVHD mice demonstrated that CD11c+ DCs exhibit a striking and significant increase beginning at day 7, peaking at day 10 and then declining through day 14 [13]. DCs play a prominent role in CD8 CTL induction, and are a major contributor to the increase in total F1 CD11c+ DC population [13]. Enhanced CTL-mediated elimination of F1 B cells in males is consistent with a more prominent role for F1 DC in CTL expansion, with greater increases in CD11c+ DCs and/or slower downregulation vs. females. We did not detect significant sex- or dose-dependent response differences in engraftment of donor B6 CD4 or CD8 T cells. Taken together, our results in acute GVHD support the conclusion that the donor anti-F1 CD8 CTL response is significantly stronger in males, with F1 B cells identified as the splenic subset most susceptible elimination.

Chronic D2→F1 GVHD is characterized by defective donor CD8 CTL SLEC maturation, resulting in continued donor CD4 T cell driven expansion of F1 splenocytes in the absence of effective CTL-mediated elimination [2, 12]. Our results demonstrate that D2 GVHD mice also exhibit a hierarchy of F1 splenic subsets targeted by donor CD4 T cells, which is distinct from B6 GVHD mice and exhibits more sex differences. Specifically, host CD4 T cells in both male and females that exhibit the greatest percentage change vs. control (males ∼280%; females 330%), rather than F1 B cells as observed in the B6 GVHD model. In males, the next highest percentage increase was observed for CD11c+ DCs (223% of control), followed by CD11b+ macrophages (189% of control) and B cells (170% of control) clustered together, and lastly CD8 T cells (118% of control). This hierarchy differed in females, with B cells exhibiting the second highest percentage increase (239%) and the remainder of the subsets clustered in the ∼175-193% range.

For both sexes, total splenocytes, F1 B cells, CD4 and CD8 T cells were all highly responsive to a small increase in donor T cells with large effect sizes (2.8-4.0) (Table 2B). Only males had a significant dose-dependent response increase in CD11b+ macrophages, and neither sex showed a significant dose-dependent increase in CD11c+ DCs, suggesting that these two myeloid subsets approached maximal expansion at the 4.5M dose. Hence F1 splenic subsets in D2 GVHD mice differ in the degree of CD4 T cell help-dependent expansion, just as they differ in their relative susceptibility to CTL lysis in B6 GVHD.

Relative to the acute model, sex differences in D2 GVHD mice were large and widespread. Females exhibited significantly greater numbers of total splenocytes, F1 CD4 T cells, F1 B cells and F1 CD8 T cells, with effect sizes ranging between 2.7-4.1 for both donor cell doses. Although total F1 CD8 T cell numbers were greater in females, males showed a much higher percentage/number of CD44^hi^ F1 CD8 T cells, consistent with more HVG activation. Previous work has shown that although the CTL response is defective in D2 GVHD mice, it is not completely absent. Both GVH and HVG CTL have been reported at day 7 in male D2 GVH mice [13]. To date, no direct male to female comparison of the relative strengths of the cytolytic GVH and HVG have been published in D2 GVHD mice. However, using donor and F1 CD8 T cell numbers as surrogate markers for the relative strength of the GVH and HVG response, males show a greater peak at day 8-10 for donor CD8 engraftment. They also exhibit greater killing of B cells at day 8, followed by earlier downregulation of donor CD4 and CD8 T cells via apoptosis and/or exhaustion [7], consistent with an intrinsic T cell affect that is androgen receptor-dependent [28–31]. The greater number of female F1 CD8 T cells at two weeks vs. males (Fig. 2D) reflects reduced downregulation in females.

Conversely, higher numbers of CD44^hi^ F1 CD8 T cells in males (Fig. 2D) is indicative of prior activation as part of a stronger male HVG reaction. By contrast, during days 8-14, female *donor-derived* T cells continue to proliferate resulting in >2-fold greater donor CD4 T cell engraftment at two weeks vs. males [5, 7]. Females also showed a higher number of both donor and F1 CD4 CD44^hi^ and CD4 Tfh cells compared to males, consistent with more robust T cell activation and skewing towards B cell help, driving F1 B cell expansion at two weeks (Fig. 2B) that persists long term [6]. Together these results support the conclusion that although the D2 SLEC CTL response is defective in both sexes, the response is transiently greater in males, as is the F1 HVG response.

Studies in humans and animals indicate that in general, females have stronger innate and adaptive immune responses than males (reviewed in [32, 33]. Following exposure to infectious pathogens or vaccinations, females generate higher antibody levels with better neutralizing capacity, consistent with lower viral burdens than males. Both female sex hormones and X-linked genes such as *TLR7* contribute to greater antibody production in females, whereas male sex hormones on the other hand dampen the immune response (reviewed in [34, 35]. This stronger antibody response also puts females at greater risk for autoimmunity, particularly for humoral autoimmune conditions such as lupus, which exhibits a 9:1 male-female ratio in disease incidence [36]. Indeed, lupus nephritis is mediated by glomerular deposition of autoantibodies, either as pre-formed circulating immune complexes or by direct binding of anti-glomerular antibodies that form in situ immune complexes (reviewed in [37]. Experimental sex hormone manipulation in NZB/W lupus mice demonstrated that female sex hormones exacerbate autoantibody-driven disease, whereas male sex hormones ameliorate disease and reduce mortality [38, 39].

Because F1 B cells are nearly completely eliminated in B6 GVHD, sex differences in humoral immunity are better revealed in D2 GVHD mice. ICGN in D2 GVHD mice is more severe in females, as evidenced by higher levels of anti-DNA and anti-PARP autoantibodies, as well as greater numbers of total splenic B cells, plasma cells and follicular B cells. [5, 7, 40]. As with NZB/W lupus mice, sex hormone manipulation in D2 GVH mice previously confirmed the role of sex hormones in ICGN severity [41]. Greater female expansion of F1 B cells observed in our study demonstrates that female skewing of ICGN disease can be detected as early as two weeks after donor cell transfer. Thus, B cell-mediated autoantibody production in the p→F1 model using BDF1 mice are in agreement with a large number of human and animal studies showing enhanced antibody responses in females.

Females are also reported to have greater CD8 CTL responses [32, 42, 43]. CD8 CTL maturation into SLEC and memory precursor effector cells (MPEC) requires not only TCR and costimulatory signals, but also a third signal such as the inflammatory cytokines IL-12 or IFNα [44]. Females are reported to exhibit stronger innate immune responses than males [45, 46], raising the question as to whether reported sex differences in CD8 CTL in infectious settings reflect greater innate immune responsiveness to inflammatory cytokines in addition to, or instead of, an intrinsic sex-based difference in CD8 CTL responses. In support for the latter idea, greater virus specific CD8 T cell responses in females following live influenza A murine infection were not seen in mice receiving an inactivated influenza A vaccine [47].

Additionally, the ability of 17β–estradiol to enhance the influenza A specific CTL response in mice was linked to recruitment of neutrophils, which can qualitatively increase the virus-specific CD8 T cell response. The CTL promoting effect of 17β–estradiol was lost in neutrophil-deleted mice [48]. Lastly, following infection with vaccinia virus or Listeria monocytogenes, greater CTL SLEC expansion is observed in female mice. This sex difference was connected to enhanced IL-12 responsiveness; when IL-12 signaling was blocked, males outpaced females in SLEC formation [49].

CD8 T cells also play a critical role in tumor immunity, where chronic antigen stimulation, rather than innate/danger signals, is the primary driver of exhaustion in mature CTL. Males have poorer overall survival rates for non-reproductive system cancers [50].

Importantly, T cell-intrinsic androgen receptor signaling functions to promote CD8 T cell exhaustion in tumor models, blunting male anti-tumor CD8 T cell activity [28–31]. These studies support the idea that males have weaker CTL responses than females even in the absence of overt innate system involvement.

Taken together, aforementioned reports highlighting enhanced CTL function in females conflict with our results in both B6 and D2 GVHD models, for which the initial in vivo SLEC response in males is as good or better than that of females. This discrepancy may relate to the distinct characteristics of the p→F1 model, in which the driving antigen is alloantigen rather than an infectious pathogen or an exogenous innate immune system activator. Also, it is not clear whether signal 3 molecules such as IL-12 and IFNα are present early on in biologically significant quantities in the p→F1 model. In preliminary studies, we were unable to detect a significant role for either of these cytokines in acute GVHD [51].

Specifically, neither transfer of B6 donor cells deficient in either IL-12R or Ifnar1, nor combined antibody-mediated blockade of both cytokines altered the acute GVHD phenotype at two weeks (manuscript in preparation). By contrast, acute GVHD could be converted to a chronic GVHD phenotype with skewing of serum Ig to IL-4 dependent isotypes (IgG1, IgE) after IL-2 blockade [52]. Blockade of TNF in the first 4 days after donor transfer also converted p→F1 mice from acute to chronic GHVD and impaired IFNg production but not production of non Th1-cytokines IL-10, IL-4 and IL-6 [53]. Similarly, transfer of B6 TNF receptor-deficient donor cells demonstrated that in the setting of a normal donor CD4 T cell IL-2 response, acute B6 GVHD was absolutely dependent on signaling through TNFR2 (p75) and not TNFR1 (p55) [20]. Together these results confirm the critical role of IL-2 in initiating acute GVHD and demonstrate that TNF, an IL-2 dependent cytokine in this context, is necessary and sufficient to drive CD8 SLEC maturation in the absence of significant levels of inducers of IFNα or IL-12. Importantly, these studies do not address a role for these signal 3 molecules in MPEC formation.

Conversely, defective donor CD8 CTL formation in D2→F1 can be corrected by administration of rIL-12 if given in the first 5 days after transfer, which converts the 2-week disease phenotype from chronic to acute GVHD [54]. Similar results were observed with early administration of CpG ODNs that induce both IL-12 and IFNα [51]

Moreover, DBA antigen presenting cells (APCs) are capable of making significant amounts of IL-12 when compared to Balb/c mice [55]. Thus, D2 mice can make IL-12 and their T cells can respond to it acutely, supporting the idea that IL-12 and IFNα are not normally induced early on in quantities sufficient to induce SLEC formation. These cytokines would be expected to substitute for TNF and promote SLEC if present early after donor cell transfer, meaning TNF blockade would not alter acute GVHD phenotype. As a caveat, these studies only examine SLEC formation over the first two weeks after donor cell infusion, and do not exclude a role for IL-12 or IFNα in the generation of memory T cell responses long term in GVHD mice.

Although lupus is notoriously heterogeneous in clinical presentation, our results have implications for pathogenesis in at least a subset of patients. Defective IL-2 production is widely reported in both human and murine lupus [56, 57]. IL-2 limits CD4 Tfh cell and GC formation [58–60] and also promotes several downregulatory mechanisms of potential importance in lupus, including formation and/or maintenance of CD8 CTL, Th1 CD4 T cells, and CD4 Tregs [61]. Defects in CD8 T cell-mediated downregulation on B cell responses have been widely reported in lupus mice [62–69]; both perforin and Fas/FasL pathways are required for optimal CD8 T cell control of B cell hyperactivity [65, 70, 71].

Whether IL-2 defects primarily predispose to lupus pathogenesis, or are secondary to disease activity, remains a major question for the field. A secondary disease-associated defect in IL-2/CTL is well-recognized (reviewed in[57]. A primary, pre-existing defect in IL-2 has been suspected, but remains difficult to address in either spontaneous murine or human lupus since it is not clear when disease is initiated. In a study of identical twins discordant for lupus, defective cytolytic CD56+ natural killer cell killing was observed in both unaffected and affected twins, consistent with a pre-existing IL-2-dependent immune defect. However, neither IL-2 production nor CTL function was directly tested [72]. The fact that D2 T cells have a major impairment in in vitro and ex vivo IL-2 production [12, 73] supports the hypothesis that a primary IL-2 defect can predispose to lupus pathogenesis. Importantly, D2 mice are not classified as immunodeficient or lupus prone - it is only when tolerance is broken in the D2→F1 setting that lupus ensues.

The pre-existing defect in IL-2 production seen in otherwise normal D2 mice supports a disease model that some lupus-predisposed individuals may have pre-existing defects in T cell IL-2 production that may go undetected, particularly in the setting of a normal innate immune response. We posit that following exposure to a lupus trigger capable of breaking tolerance in the absence of innate immune system activation, a normal lupus-resistant individual (B6-like) will respond with a strong IL-2 Th1 response that promotes active downregulatory mechanisms to eventually re-establish tolerance. Conversely, a lupus-prone individual (D2-like) exposed to the same trigger will produce less IL-2 and a weaker Th1 response, with skewing towards a Tfh response that promotes self-reactive B cell maturation and autoantibody production rather than effective downregulation or elimination. In this setting, tolerance is difficult to reestablish. In line with findings described here, females will be at greater risk for humoral autoimmunity in this scenario, due to their propensity for heightened antibody responses relative to males, coupled with a greater tendency for CD4 T cell expansion and skewing towards Tfh cells as seen in D2 GVHD mice (Figs 2C, 2H, 5E,H). Exogenous IL-2 treatment has been shown to be beneficial in lupus mice [74, 75] and human lupus (reviewed in [61]. Our results support ongoing efforts to normalize IL-2 in at least a subset of lupus patients, particularly in females.

## Acknowledgements

The authors wish to thank Cara Olsen, DrPH and Andrew Snow, PhD, USUHS for reviewing this manuscript and for helpful discussions.

## Abbreviations used

BDF1: B6D2F1
B6: C57Bl/6
D2: DBA/2
GVH: graft-vs.-host
HVG: host-vs.-graft
p→F1: parent-into-F1
MPEC: memory precursor effector cells
Tfh: T follicular helper
SLEC: short lived effector cells

